# Longitudinal Survey of Astrovirus infection in different bat species in Zimbabwe: Evidence of high genetic Astrovirus diversity

**DOI:** 10.1101/2023.04.14.536987

**Authors:** Chidoti Vimbiso, De Nys Hélène, Abdi Malika, Mashura Getrude, Pinarello Valérie, Chiweshe Ngoni, Matope Gift, Guerrini Laure, Pfukenyi Davies, Cappelle Julien, Mwandiringana Ellen, Missé Dorothée, Gori Elizabeth, Bourgarel Mathieu, Liégeois Florian

## Abstract

Astroviruses (AstVs) have been discovered in over 80 animal species including diverse bat species and avian species. A study on Astrovirus circulation and diversity in different insectivorous and frugivorous chiropteran species roosting in trees, caves and building basements was carried out at 11 different sites across Zimbabwe. Pooled and individual faecal sampling methods were used for this study, with collection dates ranging from June 2016 to July 2021. In two sites, Magweto and Chirundu, sampling was carried out at monthly intervals from August 2020 to July 2021. Astroviruses and bat mitochondrial genes were amplified using pan-AstVs and CytB /12S RNA PCR systems respectively. Phylogenetic analysis of the *RdRp* Astrovirus sequences revealed a high genetic diversity of astroviruses. All the bat astroviruses tested in this study clustered with the *Mamastrovirus* genus. Two distinct groups of amplified sequences were identified. One group was comprised of sequences isolated from *Hipposideros, Rhinolophus* and *Eidolon helvum* spp. clustered with Human Astrovirus strains: *HuAstV* types1-6, *HuAstV*-MLB1-3 and *HuAstV*-VA-1. The second group comprising the majority of the sequences clustered with different strains of Bat AstVs. Results from the longitudinal study at Magweto and Chirundu showed an overall AstV prevalence of 13.7% and 10.4% respectively. Peaks of AstV infection at Chirundu coincided with the period when juveniles are 4-6 months old. By combining data from our previous work on Coronaviruses, we also anaylzed co-infection of bats with Coronaviruses and Astroviruses at Magweto and Chirundu sites for which the prevalence of co-infection was 2.6% and 3.5% respectively.

## Introduction

Astroviruses (AstVs) are non-enveloped single-stranded positive sense RNA viruses with a genome length of approximately 6.2 to 7.7 Kb (<u>Bosch et al., 2014</u>). They have been described in over 80 non-human species and are classified into two genera: *Mamastrovirus* and *Avastrovirus,* respectively infecting mammals and avian species (<u>Donato & Vijaykrishna,</u> <u>2017</u>). According to the International Committee on Taxonomy of Viruses (https://talk.ictvonline.org), *Mamastrovirus* are classified into 19 recognized species, *MAstV*-1 to –19, and two genogroups GI and GII (ICTV, 2020). All classic and novel Human Astrovirus (*HuAstVs*) belong to four different species, *MAstV*-1, *MAstV*-6, *MAstV*-8 and *MAstV*-9 <u>(Boujon</u> <u>et al., 2017</u>; <u>Donato & Vijaykrishna, 2017</u>; <u>Wohlgemuth et al., 2019</u>). Recently, studies have reported AstVs infections in fish and insects (<u>Wu et al., 2020</u>; <u>Roach & Langlois, 2021</u>), however bats and wild birds are considered to be natural reservoirs of the *Astroviridae* family (<u>Fischer et al., 2016</u>; <u>El Taweel et al., 2020</u>). Generally *HuAstVs* have been identified as causal agents of acute viral gastrointestinal illness worldwide particularly in children, immunocompromised people and the elderly (<u>El Taweel et al., 2020</u>; <u>Wu et al., 2020</u>; <u>Roach &</u> <u>Langlois, 2021)</u>. Of note, in humans, the majority of AstV-associated encephalitis or meningitis illnesses were reported in immunocompromised people (<u>Frémond et al., 2015</u>; <u>Cortez et al.,</u> <u>2017</u>; <u>Janowski et al., 2019</u>). Furthermore, beyond this well-known clinical manifestation of AstVs infections, neurovirulent AstVs infections have also been reported in both humans and domestic animals (<u>Vu et al., 2016</u>; <u>Johnson et al., 2020</u>). Moreover, studies have reported numerous cross-species transmissions of AstVs (<u>Roach & Langlois, 2021</u>), where some AstVs detected in humans closely clustered with strains identified in rodents, minks, feline species and ovine (<u>Donato & Vijaykrishna, 2017</u>; <u>Wohlgemuth et al., 2019</u>; <u>Roach & Langlois, 2021</u>). Bat-borne viruses represent an extensive research field owing to the plethora of viruses hosted by Chiropterans (<u>Baker et al., 2013</u>). Furthermore, this mammalian order is known to be persistently infected by astroviruses (<u>Chu et al., 2008</u>) and this is inclusive of many species of insectivorous bats (<u>Dufkova et al., 2015</u>). To date, a high diversity of bat AstVs from different bat families has been reported worldwide (<u>Rougeron et al., 2016</u>; <u>Cortez et al., 2017</u>; <u>Fischer</u> <u>et al., 2017</u>). Bat AstVs belong to *MAstV*-12, and *MAstV*-14 to –19 species (<u>Bosch et al., 2014</u>). In Europe a variety of bat species belonging to the Yangochiroptera sub-order have been discovered to harbor several astroviruses (<u>Lacroix et al., 2017</u>). In Africa, bat astroviruses were reported in Egypt, Gabon, Madagascar, Kenya and Mozambique (<u>Rougeron et al., 2016</u>; <u>Waruhiu et al., 2017</u>; <u>Lebarbenchon et al., 2017</u>; <u>Hoarau et al., 2018</u>).

AstVs are known to occur in bat species with high prevalence and exceedingly high genetic diversity whereby infections in these hosts are usually non-pathogenic (<u>Lee et al., 2018</u>; <u>Bergner et al., 2021</u>). Despite all these studies, large gaps exist on astroviruses prevalence and infection dynamics in bats (<u>Fischer et al., 2016</u>). Astroviruses prevalence and detection in bat colonies seemed rather correlated to abiotic factors such as seasons and year of sampling and biotic factors such as sex and reproductive status and also viral co-infection (<u>Seltmann et al.,</u> <u>2017</u>). Viral shedding in bats occurs in spatial and temporal pulses that can drive spillover to other animals or humans (<u>Drexler et al., 2011</u>; <u>Plowright et al., 2016</u>). The varying prevalence of viruses in bats is determined by shedding pulses, where rare or very low prevalences are detected in the absence of shedding pulses and higher prevalence is detected when shedding pulses occur (<u>Amman et al., 2014</u>; <u>Plowright et al., 2016</u>).

Until now, no data was available on genetic diversity and prevalence of circulating AstVs in insectivorous bats in Zimbabwe. In this study we enlarge the spectrum of bat AstVs knowledge in Africa by identifying and analyzing bat-AstVs, their diversity and prevalence from different bat species colonies according to reproductive season phenology in Zimbabwe. Furthermore, we will also show and highlight the presence of co-infection of bats by astroviruses and coronaviruses (CoVs) and how it varies according to reproductive phenology.

## Material and Methods

### Sampling approaches and sites

Two different approaches were followed in this study: bat community sampling and individual bat sampling.

### Bat community pooled sampling

Between February 2016 and December 2020, faecal samples from both insectivorous and frugivorous bat species were respectively collected in different sites including caves, an ancient mine and trees in Zimbabwe **(Figure 1**, **Table 1)**.

**Figure 1:**
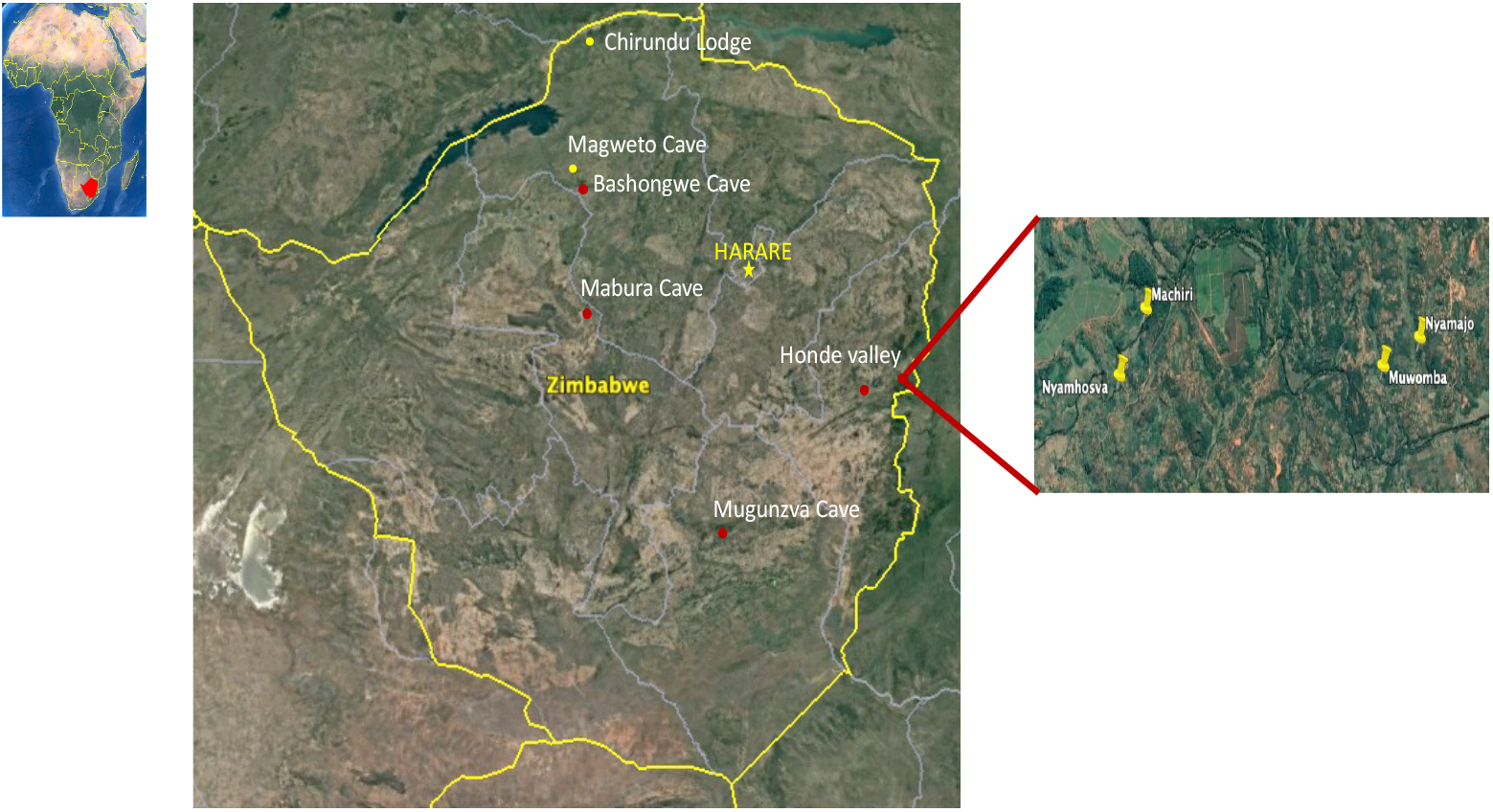
The study sites highlighted on the map show the sampling areas for insectivorous and frugivorous bats colonies across Zimbabwe. In Honde valley, the *Eidolon helvum* colony was established in at least four different sites represented by the rectangle at the right of the figure. Red circles represent community approach study sites; yellow circles represent longitudinal survey sites (individual approach).

**Table 1:** Number of Astrovirus positives per site/sampling year in pooled faecal samples from frugivorous and insectivorous bats for the selected 14 sites.

| Sampling Sites | Type of roost | Collection Date | Bat Species | Number of collected tubes | Number of Tested pools | Number of positive pools |
| --- | --- | --- | --- | --- | --- | --- |
| Magweto Cave | Cave | June 2016 | *Hipposideros spp.* | 24 | 8 | 5 |
| Bashongwe Cave | Cave | March 2018 | *Rhinolophus and Hipposideros spp.* | 59 | 7 | 0 |
|  |  | July 2018 |  | 57 | 20 | 0 |
|  |  | December 2018 |  | 135 | 45 | 0 |
| Mabura Cave | Cave | June 2016 | *Hipposideros spp.* | 36 | 12 | 6 |
|  |  | February 2017 | *Hipposideros spp.* | 22 | 7 | 0 |
| Mugunza Cave | Cave | April 2017 | *Roussetus aegyptus* | 50 | 13 | 0 |
| Honde Valley | Trees |  |  |  |  |  |
| Machiri |  | August 2017 | *Eidolon helvum* | 25 | 9 | 0 |
| Muwomba |  | November 2017 | *Eidolon helvum* | 66 | 22 | 0 |
| Nyamajo |  | November 2017 | *Eidolon helvum* | 58 | 19 | 0 |
| Nyamhosva |  | November 2017 | *Eidolon helvum* | 35 | 11 | 0 |
| Nyamajo |  | January 2018 | *Eidolon helvum* | 17 | 6 | 1 |
| Machiri |  | January 2018 | *Eidolon helvum* | 7 | 2 | 2 |
| Muwomba |  | January 2018 | *Eidolon helvum* | 8 | 2 | 0 |
| Nyamhosva |  | January 2018 | *Eidolon helvum* | 28 | 10 | 1 |
| Nyamajo |  | February 2020 | *Eidolon helvum* | 92 | 92 | 0 |
| Chirundu | Tree | December 2020 | *Rhinolophus spp.* | 21 | 21 | 1 |
| **Total** |  |  |  | **740** | **306** | **16** |

All these sites were chosen according to the presence of bat colonies and the existence of anthropic activities. For instance, the selected caves and ancient mine are regularly visited by local people to collect bat guano, which is used as fertilizer, and/or to hunt bats for consumption. In Honde valley, frugivorous bats roosting sites in trees were close to maize crops or/and orchards.

All sites, except for three, were visited at different times during the sampling period **(Table 1)**. The same sampling method was used at all sites and in every session as previously described by Bourgarel et al, (<u>Bourgarel et al., 2018</u>). Briefly, two square meters of plastic sheets were laid down at each site/per sampling session, underneath the bat colonies and left overnight. Six grams of fresh faeces from different bat individuals were collected from each plastic sheet in a 15 ml tube with 6 ml of RNA stabilization solution (https://protocol-online.org*/*). Back in the laboratory, samples were stored at –80°C for subsequent analysis. Of note, for all sampling sessions, personnel protection equipment (double pair of gloves, FFP3 mask, Tyvek overall, goggles, boots) were used in order to respect the biosafety procedures and to protect the sampling teams. Good biosafety practices training are regularly dispensed for our sampling teams.

### Bat individual sampling

Individual bat samples which had already been collected from two study sites for a study on coronaviruses by Chidoti et *al*. were used in this study (Figure 1) (<u>Chidoti et al., 2022</u>). These two sites, one cave (Magweto) and one building basement (Chirundu Farm) had been visited from August 2021 to July 2022 on a monthly basis **(Figure 1**, **Table 2)** (<u>Chidoti et al.,</u> <u>2022</u>). Unfortunately, this study was conducted during the COVID 19 crisis and we were not able to access the study sites every month as planned owing to the imposed lock downs. Faecal samples had been collected by placing two square meters plastic sheets underneath the bat colonies. Only one faecal dropping (not contaminated by other faeces or urine) per 20 cm^2^ was collected, assuming it represented one individual. Faeces were conserved individually in a 1.5 ml tube filled up with 0.5 ml of home-made RNA stabilization solution (https://protocol-online.org*<u>/</u>*) and stored at –80°C before further laboratory analyses.

**Table 2:** Prevalence of Astroviruses and co-infection with Coronaviruses and confidence intervals (CIs) per month at both Magweto and Chirundu sites in faecal samples from individual sampling of insectivorous bat communities and per reproduction cycle stage.

| Site | Reproduction cycle | Month sampled | No of samples tested | No of Astrovirus positives | Prevalence (%) + CI (95%) | No of coinfection CoV and AstV | Prevalence (%) + CI (95%) |
| --- | --- | --- | --- | --- | --- | --- | --- |
| Chirundu | Non-gestation | August, 2020 | 153 | 0 | 0.0 (0.0-0.2) | 0 | 0.0 (0.0-0.2) |
| Chirundu | Pregnancy | October, 2020 | 297 | 3 | 1.0 (0.3-2.9) | 0 | 0.0 (0.0-1.3) |
| Chirundu | Parturition | November, 2020 | 241 | 3 | 1.2 (0.4-3.6) | 1 | 0.4 (0.05-2.3) |
| Chirundu | Lactation | December, 2020 | 159 | 16 | 10.1 (6.3-15.7) | 7 | 4.4 (2.1-8.8) |
| Chirundu | Weaning | February, 2021 | 170 | 29 | 17.1 (12.1-23.4) | 18 | 10.6 (6.8-16.1) |
| Chirundu | 4-6 Month juveniles | March, 2021 | 240 | 51 | 21.3 (16.5-26.8) | 19 | 7.9 (5.1-12.0) |
| Chirundu | Non-gestation | May, 2021 | 243 | 47 | 19.3 (14.8-24.7) | 9 | 3.7 (1.9-6.8) |
| Chirundu | Non-gestation | July, 2021 | 225 | 30 | 13.3 (9.5-18.4) | 9 | 4.0 (2.1-7.4) |
| Overall prevalence |  |  | 1728 | 179 | 10.4 (9.0-11.8) | 63 | 3.6 (3.8-5.8) |
| Magweto | Non-gestation | September, 2020 | 348 | 18 | 5.2 (3.3-8.0) | 1 | 0.3 (0.05-1.6) |
| Magweto | Pregnancy | October, 2020 | 257 | 46 | 17.9 (13.7-23.0) | 7 | 2.7 (1.3-5.5) |
| Magweto | Parturition | November, 2020 | 228 | 31 | 13.6 (9.7-18.6) | 3 | 1.3 (0.4-3.8) |
| Magweto | 4-6 Month juveniles | March, 2021 | 242 | 46 | 19.0 (14.5-24.4) | 20 | 8.3 (5.4-12.4) |
| Magweto | Non-gestation | April, 2021 | 242 | 26 | 10.7 (7.4-15.2) | 6 | 2.5 (1.1-5.3) |
| Magweto | Non-gestation | June, 2021 | 242 | 47 | 19.4 (14.9-24.9) | 3 | 1.2 (0.4-3.6) |
| Overall prevalence |  |  | 1559 | 214 | 13.7 (12.1-15.5) | 40 | 2.6 (1.8-3.5) |

Periods of gestation (pregnancy), parturition, lactation, weaning and presence of 4-6 months old juveniles had been determined based on observations (from captures and observation of the roosting bats) combined with literature, as already reported in Chidoti et *al.* (<u>Chidoti et al., 2022</u>). The reproductive season at Chirundu and Magweto site had been observed to begin in September to February for the predominant bat species. The different insectivorous bat families observed at both sites had been found to be synchronous regarding their reproductive cycles, and consensus reproduction periods had been determined based on the literature and observations (<u>Monadjem et al., 2010</u>; <u>Chidoti et al., 2022</u>).

## Nucleic acid extraction and RT-PCR

### Community samples

Nucleic acids extraction was done from all faecal samples as previously described [30]. Briefly, biological material (faeces) of three or four sample tubes collected from the same plastic sheet were pooled and transferred in a 50 ml tube with 20 ml of PBS 1X then vigorously mixed. Tubes were centrifuged at 4500 rpm for 10 min. The supernatant was first filtered using gauze in order to eliminate faecal matter and transferred in fresh tubes then re-centrifuged at 4500 rpm for 10 min. The supernatant was filtered through a 0.45 µm filter to remove eukaryotic and bacterial sized particles. Seven millilitres of filtered samples were centrifuged at 250,000 g for 2.5 h at 4°C. The pellets were re-suspended in 600 µl H_2_0 molecular grade and 150 µl were used to extract RNA and DNA using NucleoSpin® RNA Kit (Macherey-Nagel, Hoerdt, France) according to the manufacturer’s protocol.

### Individual samples

Nucleic acids were extracted from 200 μl of faecal samples preserved in 0.5 ml RNA stabilization solution using 5X MagMax Pathogen RNA/DNA Kit (ThermoFisher Scientific, Illkirch-Graffenstaden, France), as already described in (Bourgarel et al., 2018) The faeces were vortexed vigorously (30Hz) using Retsch MM400 Tissuelyser for 5 min to fully homogenise and mix the faecal particles, followed by centrifugation at 16000 g for 3 min to fully separate the supernatant from the faecal debris. A volume of 130μl of the supernatant was used for the isolation and purification stage of the nucleic acids using Mag Max extraction kit with the automatic KingFisher Duo Prime Purification System extractor (ThermoFisher Scientific, Illkirch-Graffenstaden, France). A final volume of 80μl of eluted RNA/DNA was stored at –80◦C.

### Bat species identification

For all bat samples that tested positive for AstVs, bat species identification was carried out by the amplification and sequencing of mitochondrial C*ytochrome b* gene as previously described by Kocher et *al.* (<u>Kocher et al., 1989</u>). *S*equences were then compared to available bat sequences in the GenBank database using *Basic Local Alignment Search Tool* (BLAST).

### Astrovirus detection

Reverse Transcription (RT) using random hexamers were done on 5μl of RNA sample template using 1μl random hexamers, 0.5μl Oligo dT primer, 0.4μl of dNTPs (10mM) (ThermoFisher Scientific, Illkirch-Graffenstaden, France) and 5.5μl molecular grade water incubated at 65°C for 5 min. This was followed by addition of 4μl of Buffer 5X, 2μl of 0.1M DTT (M-MLV Reverse Transcriptase, Invitrogen, ThermoFisher Scientific) and 1μl of RNAse OUT, incubated at 37°C for 2 min. A volume of 1μl of M-MLV (M-MLV Reverse Transcriptase, Invitrogen, ThermoFisher Scientific, Illkirch-Graffenstaden, France) reverse transcriptase was added to the mixture followed by incubation at 25°C for 10 min, 37°C for 50 min and 70°C for 15 min. The cDNA obtained was then used to partially amplify the *Astrovirus RNA-dependent-RNA polymerase* gene (*RdRp*) by using a semi-nested Pan-Astrovirus PCR system developed by Chu et *al* (Chu et al., 2008). Visualization of positive PCR products was performed with agarose gel electrophoresis using SYBR green on a 1% gel. Positive PCR products (422 bp) were gel-agarose or directly purified (Geneclean Turbo Kit, MP Biomedicals, Illkirch-Graffenstaden, France) and then sequenced in both 5’ and 3’ directions (LGC, Berlin, Germany) by using Sanger method.

For the community approach, purified PCR products were cloned by using Topo PCR Cloning kit according to the manufacturer’s protocol (ThermoFisher Scientific, Illkirch-Graffenstaden, France). Ten clones per PCR product were sequenced in both 5’ and 3’ direction using the Sanger method (Eurofins, Germany).

## Phylogenetic analyses

Bat AstVs overlapping sequences were assembled into contiguous sequences using Geneious software package V. 2021.2.2 (Biomatters Ltd, Auckland, New Zealand). Partial non-concatenated nucleic acid sequences of the new AstVs were aligned using Clustal W (<u>Larkin</u> <u>et al., 2007</u>) implemented in MEGA 7 (<u>Kumar et al., 2016</u>), with minor manual adjustments. Ambiguously aligned and divergent regions were excluded from subsequent analyses. Phylogenies were inferred using Maximum Likelihood (ML) method implemented in PhyML V. 3.0 (<u>Guindon et al., 2010</u>). The suited evolution model was selected by Akaike’s Information criterion (AIC) using Topali software (<u>Milne et al., 2009</u>). The reliability of branching orders was tested using the bootstrap approach (1000 replicates) (<u>Lemoine et al., 2018</u>) and the GTR + F+ I substitution model was determined as the best suited evolution model.

## Temporal variations of Astrovirus prevalence and bat reproductive phenology

In the same way as in Chidoti et *al*, the prevalence of astrovirus infection was calculated at the community level for small insectivorous bat species from the two longitudinal study sites, Magweto and Chirundu farm (<u>Chidoti et al., 2022</u>). The proportion of RNA AstV-positive samples were estimated per month and site with 95% confidence intervals (CI) using Wilson score test (https://epitools.ausvet.com.au/ciproportion) (<u>Wilson, 1927</u>).

The influence of the different phases of the bat reproductive cycle stages (periods of pregnancy, parturition/lactation, weaning, weaned juveniles of 4 to 6 months old) and of *Coronavirus* infection/shedding on the prevalence of astroviruses was tested by running a generalized linear mixed model (GLMM) for each site (Magweto and Chirundu) as described in Chidoti et *al* (<u>Chidoti et al., 2022</u>). Parturition and lactation were analysed as one season because the observations did not allow the separation of the two periods as they were interlinked. The response variable with a binomial distribution was the AstVs PCR result of the samples, and the explanatory variables with fixed effects were the different phases of the reproduction cycle. Given Coronavirus PCR results on the same samples were available through our previous study (see below), Coronavirus status was also integrated into the model as fixed variable to investigate any effect of co-infection on AstVs infection/shedding. The reproductive phases were coded as 1 if it was during the corresponding reproduction phase and 0 if it was not. A session identification code was included as a random effect to control for repeated measures from the same sampling session to account for clustered samples collected.

### Astrovirus and Coronavirus co-infection

Coronavirus data, produced by Chidoti et *al* (<u>Chidoti et al., 2022</u>), on the same sample sets were used in this study to assess the AstVs and CoVs co-infection in bat communities from Magweto and Chirundu sites and a descriptive analysis of prevalences was carried out (**Table 2**).

## Results

### Astrovirus detection in pooled sampling sites

The sampling was carried out from June 2016 to December 2020 **(Table 1)** at the selected sites. At Magweto cave site, 24 samples were collected. In Bashongwe collection was carried out in March, July and December 2018, and 251 samples were collected. For Mabura cave site, a total of 58 samples were collected, in 2016 and 2017, with sampling only carried out once in both years. In Mugunza cave site, a once off sampling session was done in 2017 and a total of 50 samples were collected. In the Honde Valley site, 336 samples were collected under nine *Eidolon helvum* roost trees (Table 1). In Magweto, five of the eight pooled samples (62.5%) tested positive for AstV. In Bashongwe cave, all the 72 samples collected tested negative for AstV **(Table 1)**. In Mabura, Six (31.6%) out of 19 pooled samples tested positive for AstVs and all of the 13 samples from Mugunza tested for AstVs were negative (Table 1). In Honde Valley, 173 pooled samples were tested for AstVs and only four (2.3%) were positive **(Table 1)**. In Chirundu Baobab tree only one (4.8%) out of the 21 samples collected was positive for AstV.

### Prevalence and seasonality of Astroviruses at Chirundu and Magweto site

In total, we analyzed 1559 samples from Magweto cave site, and 1728 samples from Chirundu farm. (Table 2). The overall prevalence of AstVs at bat community level at Chirundu was 10.4% [95% CI: 9.0-11.8] (179 positives out of the 1728 samples). The highest prevalence of 21.3% [95% CI: 16.5-26.8] was observed in March 2021 and corresponded to the 4-6 months old juvenile period (Table 2). The prevalence of AstV increased from December 2020 to March 2021, which corresponded to the lactation period, followed by the weaning of four to six months old juveniles. The lowest prevalence was observed in August (0%), October (1% [95% CI: 0.3-2.9]) and November (1.2% [95% CI: 0.4-3.6]), which correspond to the transition between non-gestation and gestation period, as well as the beginning of the parturition period **(Table 1**, **Figure 2a)**.

**Figure 2:**
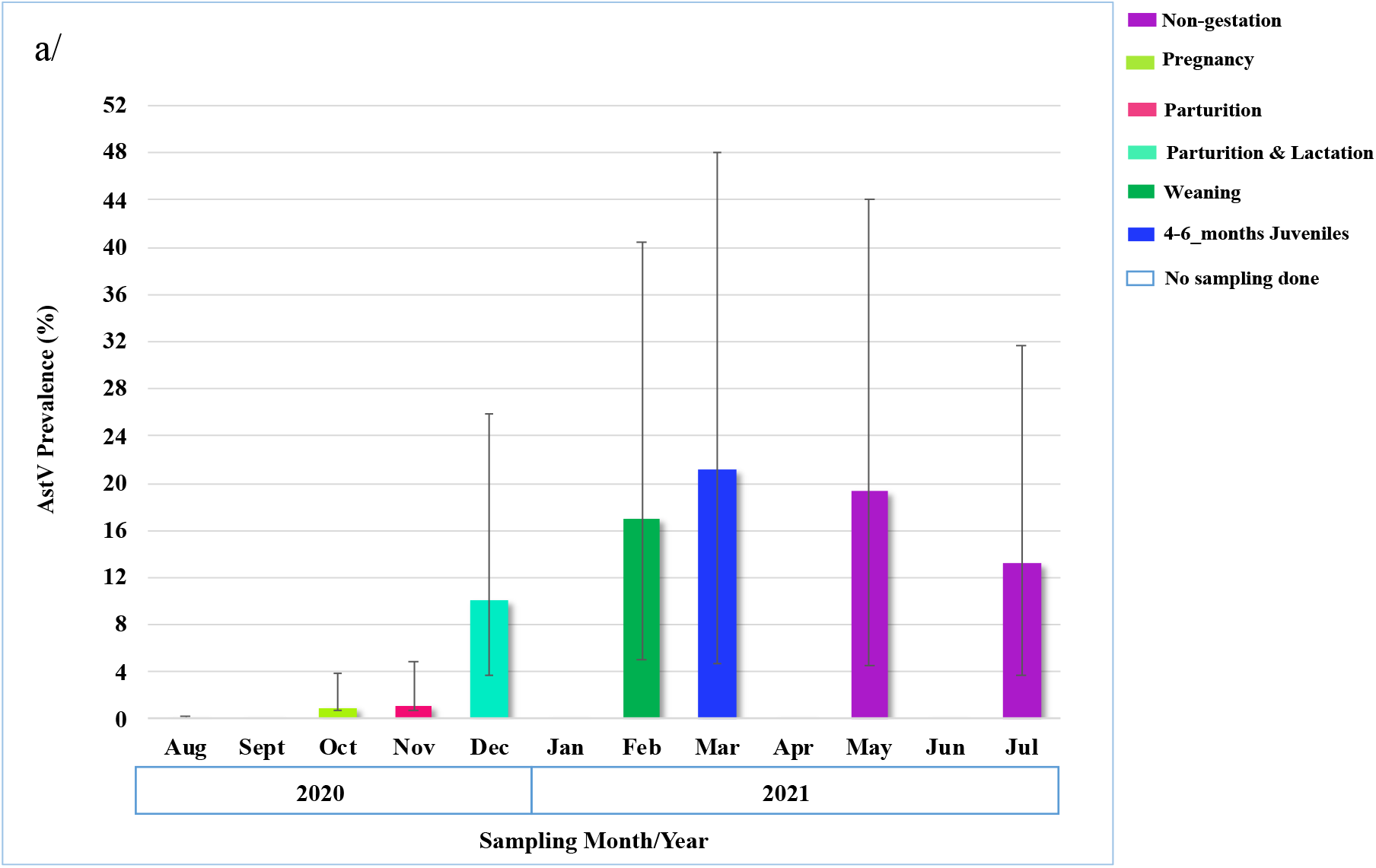
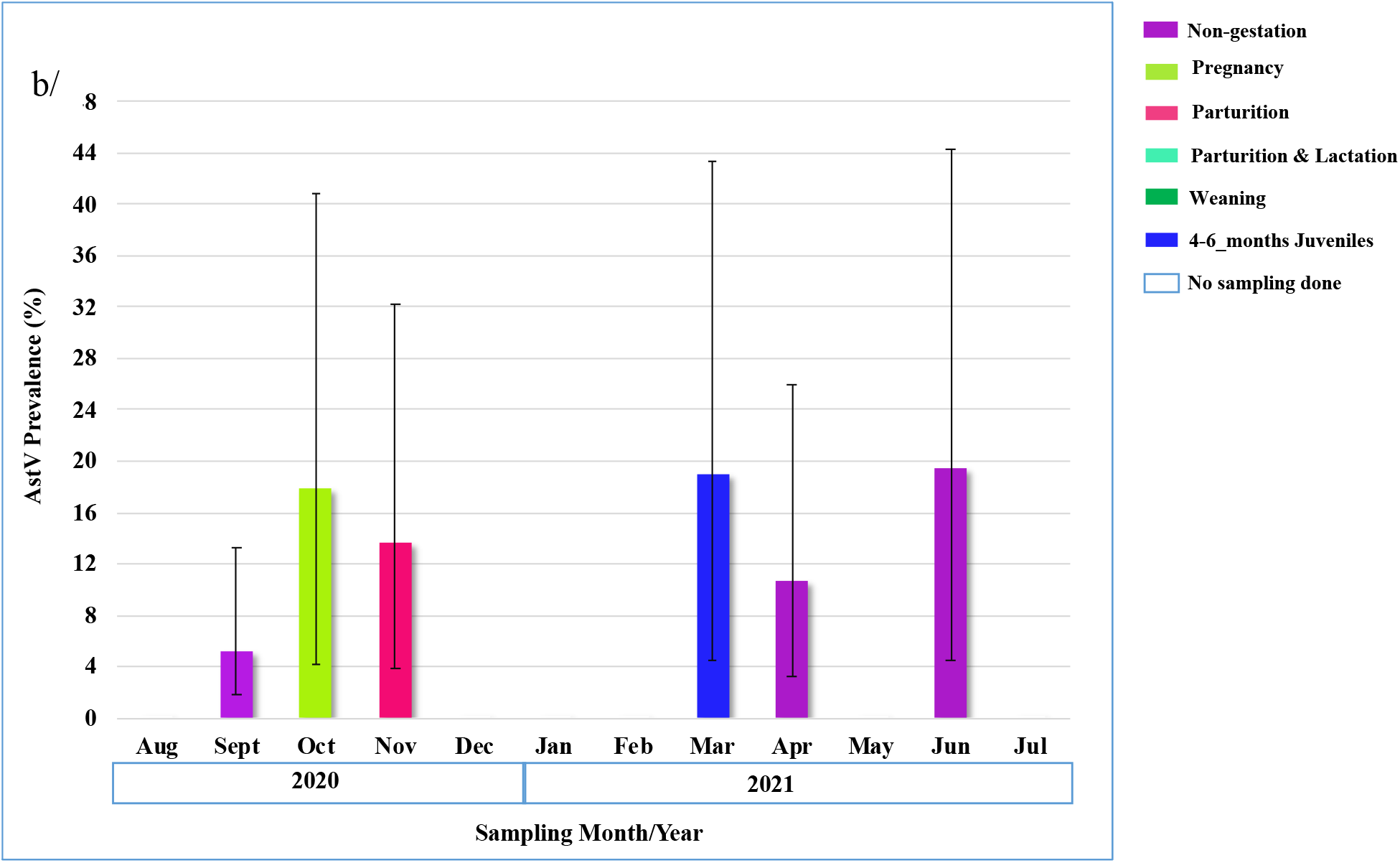
Bat community AstVs prevalence per month at Magweto site. **Fig 2a & 2b** show estimation of the astrovirus prevalence (with CI 95%) at Chirundu (a) and Magweto (b) sites respectively for the months and year sampled. The graphs were plotted with y-axes as the percentage of AstVs prevalence and x-axis as the sampling collection date and the corresponding reproductive cycle stage, each colour code representing a different season.

In Magweto site, the overall prevalence of Astroviruses for the insectivorous bat community was 13.7% [95% CI:12.1-15.5] (214 positives out of 1559 samples). There was no clear pattern for the prevalence of AstVs according to reproductive cycle stages in this site **(Table 2**, **Figure 2b)**. The highest prevalence of 19.4% [95% CI: 14.9-24.9] was observed in June 2021 during the non-gestation period, and very close prevalences observed during the pregnancy (17.9% [95% CI: 13.7-23]) and 4-6 months old juvenile (19% [95% CI: 14.5-24.4]) periods **(Table 2)**. Lower prevalence was observed in September at the end of the non-gestation period (5.2% [95% CI: 3.3-8]).

Results from the GLMM didn’t show any effect of bats seasonal reproduction periods on detection of AstV RNA in samples collected at Magweto site **(Table 3)**. For Chirundu site, the GLMM showed significantly higher probability of being positive to astrovirus during the period of weaned 4-6 months old juveniles (odds ratio (OR) = 5.088, 95% CI = 1.33-36.32, p= 0.016), and significantly lower probability of AstV positivity during the pregnancy period (odds ratio (OR) = 0.136, 95% CI = 0.03-0.81, p=0.006) **(Table 3)**.

**Table 3:** Results of the GLMM with the following explanatory variables for both site (Chirundu and Magweto): PCR_Pan-*Coronavirus*, Pregnancy, Weaning, Weaned juveniles 4 to 6 months. With references OR odds ratio, CI95 95% confidence interval.

| Variable | Odds ratios | p-value | OR CI95 |
| --- | --- | --- | --- |
| **Chirundu Site** |  |  |  |
| Intercept | 0.05 | 6.68e-09 | 0.010-0.121 |
| PCR Pancoronavirus | 1.116 | 0.536 | 0.785-1.574 |
| Pregnancy | 0.14 | 0.006 | 0.27-0.810 |
| Weaning | 3.91 | 0.094 | 0.721-39.593 |
| Weaning Juveniles 4 to 6 months | 5.09 | 0.016 | 1.333-36.322 |
| **Magweto Site** |  |  |  |
| Intercept | 0.235 | 0.0005 | 0.088-0.627 |
| PCR Pancoronavirus | 1.238 | 0.290 | 0.825-1.825 |
| Pregnancy | 0.473 | 0.145 | 0.138-1.548 |
| Weaning Juveniles 4 to 6 months | 0.678 | 0.448 | 0.202-2.256 |

### Prevalence and seasonality of coinfection with Astroviruses and Coronavirus at Chirundu and Magweto site

The co-infection prevalence in bat communities from Chirundu and Magweto sites was relatively low. The co-infection prevalence was observed to follow a similar trend as that of CoVs and AstVs prevalences, where it showed peaks that correlated with peaks of the individual viral families **(Figure 3a & b**). The overall prevalence of co-infection at Chirundu site was 3.6% [95% CI: 3.8-5.8] (Table 2) with the highest co-infection prevalence of 10.6% [95% CI: 6.8-16.1] observed in February 2021 during the weaning period. The lowest prevalences of 0% were observed in August [95% CI: 0.0-0.2] and October [95% CI: 0.0-1.3] during the end of the non-gestation and the pregnancy period, respectively **(Table 2**, **Figure 3a)**.

**Figure 3:**
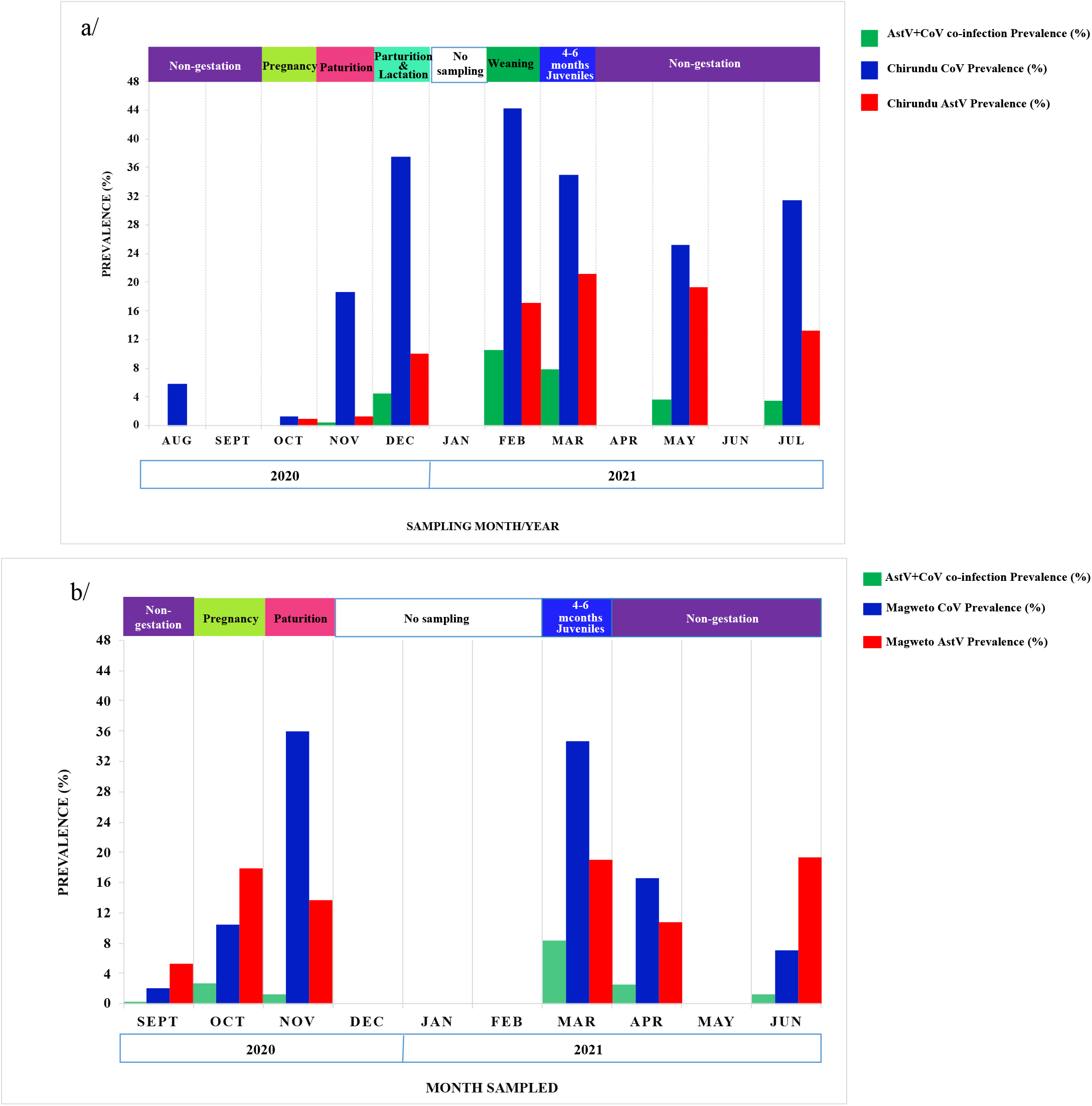
Prevalence of *Astrovirus*, *Coronavirus* and co-infections involving both viruses per month: co-infections are represented in green, coronaviruses in blue and astroviruses in red for a/ Chirundu site and b/ Magweto site according to reproduction cycle stages.

The overall prevalence of co-infection at Magweto site was 2.6% [95% CI: 1.8-3.5] **(Table 2**, **Figure 3b)**. Co-infection in this site was recorded from September 2020 to December 2020 and from March 2021 to June 2021 **(Table 2**, **Figure 3b)**. The highest prevalence of coinfection was 8.3% [95% CI: 5.4-12.4] in March during the 4-6 months old juvenile period; while the lowest 0.3% [95% CI: 0.05-1.6] was observed in September 2020 during the non-gestation period **(Table 2**, **Figure 3b)**.

### Astroviruses genetic diversity

All AstV positive samples (N=423) were sequenced, representing 179 samples from Chirundu, 229 from Magweto, seven from Mabura, seven from Honde valley, and one from Baobab tree. Of the 423 samples, 214 (50.6%) were amplified from *Hipposideros* spp., 63 (14.9%) from *Rhinolophus* spp., 53 (12.5%) from *Miniopterus* spp., 10 (2.4%) from *Nycteris* spp., seven (1.7%) from *Eidolon* spp. and for the remaining 76 samples known to originate from bats for which we couldn’t determine the genus/species.

A phylogenetic analysis was conducted with 155 partial sequences generated during this study and representative of the genetic diversity observed. All clustered within the *Mamastrovirus* genus **(Figure 4)**.

**Figure 4:**
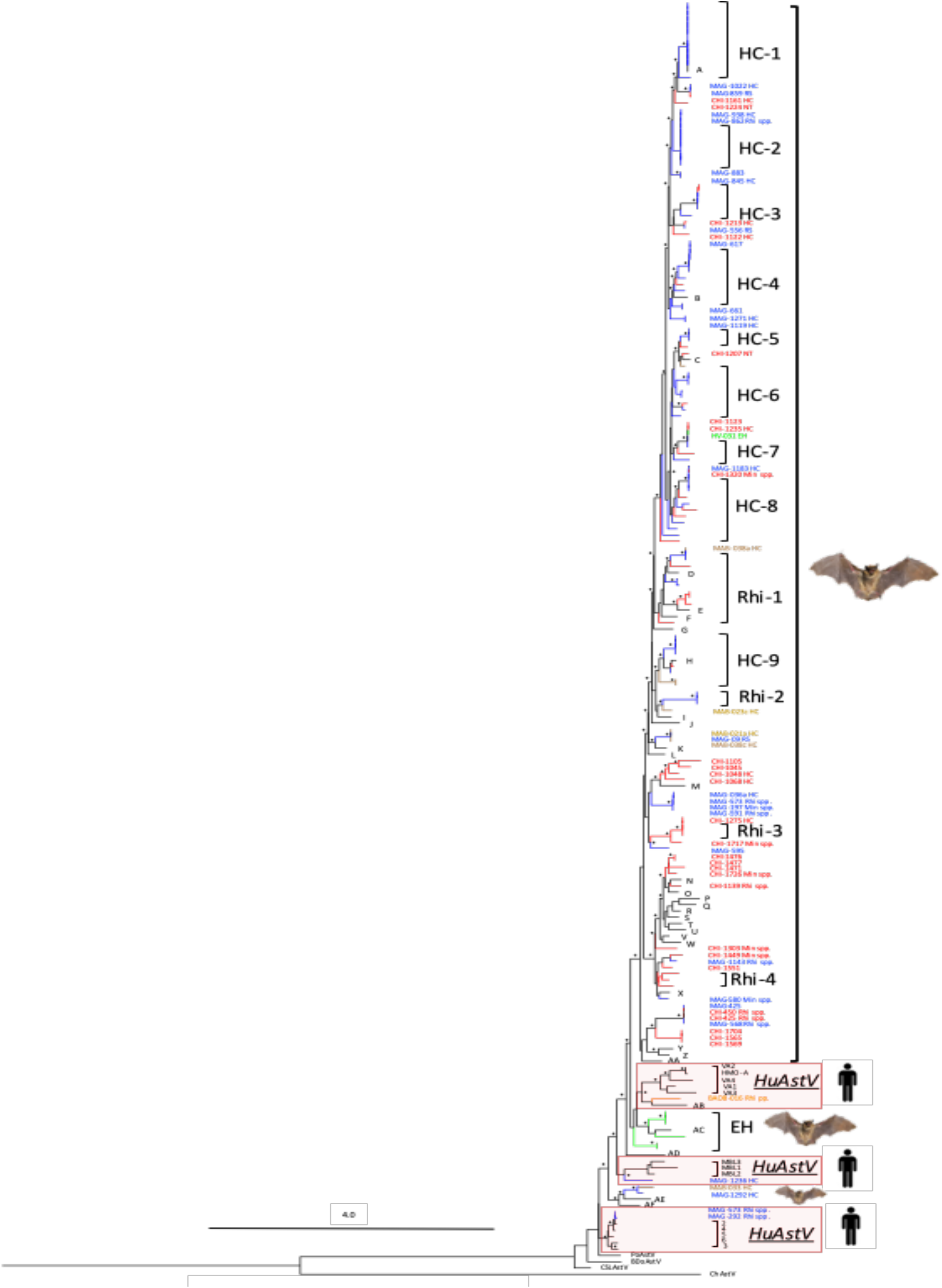
Phylogenetic tree of *Astroviruses* partial *RdRp* gene. The sequences detected at Chirundu site are represented by red branches, at Magweto site by blue branches, at Honde Valley by green branches, at Mabura cave by brown branches and at Baobab tree site by an orange branch. Human *Astrovirus* are highlighted in red rectangles and bold letters represent the different bat AstV references used to build this tree (See Supplementary Data: Figure 4). The tree was built using the maximum likelihood method based on the GTR + G4 + I model. The robustness of nodes was assessed with 1000 bootstrap replicates. Bootstrap values >70 are represented by asterisks and those <70 are not shown. HC=*Hipposideros caffer*; RS=*Rhinolophus simulator*; NT=*Nycteris thebaica*; EH= *Eidolon helvum*; Min=*Miniopterus* spp.; Rhi=*Rhinolophus* spp.. MAG=Magweto site; CHI=Chirundu site; BAOB= Baobab; MAB= Mabura site; HV=Honde valley site. The scale bar represented the number of substitutions per site. The AstV sequences used as references are detailed in supplementary data: DOI **10.5281/zenodo.8399361**

Two sequences isolated from Magweto cave, one from *Rhinolophus* spp. (MAG-573) and 1 from *Hipposideros* spp. (MAG-292) clustered with a group of Human astroviruses (*HuAstVs*) **(Figure 4)**. This cluster was well supported (Bootstrap >90%) and the nucleotide acid identities between Bat Astroviruses MAG-573, MAG-292 and *HuAstV*-2 were 96 and 93% respectively (Data not shown).

The other specific cluster of bat astrovirus showing a close relation to *HuAstVs* is a sequence isolated from *Hipposideros* spp. (MAG-1236) in Magweto, it clustered with *HuAstV-MLB-1, –2* and *3* **(Figure 4)**. Six sequences isolated from the frugivorous bats *E. helvum* found in Honde Valley and one from *Rhinolophus* spp. collected in a Baobab tree formed a sister clade with another group of Human astroviruses *HuAstV-VA-1, –2, –3, –4* and *HuAstV-HMO-A* (Figure 4). In this study, the majority of the sequences were closely related to astroviruses identified in insectivorous bats **(Figure 4)**. These were mainly amplified from *Hipposideros* spp. (HC-1 to –9) and *Rhinolophus* spp. (Rhi-1 to –4) bat species. One cluster comprised specifically of sequences isolated from *Rhinolophus* spp. Most of these sequences were isolated from Chirundu and a few of them from Magweto, and clustered together with AstV strains from *Nyctalus* spp. and *Verpitillio* spp. bat species isolated in Czech Republic (references Y & Z in the phylogenetic tree) (Figure 4). Another cluster comprising of the majority of AstV sequences from Chirundu and some from Magweto site, were characterized from *Miniopterus* spp. and *Rhinolopus* spp. All together, they formed a phylogenetic cluster with strains of bat astroviruses isolated from *Miniopterus* spp. from Madagascar and China (references N, O, R, U, T), *Rhinolophus* spp. from Korea (references P, Q) as well as *Rousettus* spp. and *Paratrianops* spp. Bat AstVs *(BAstV)* from Madagascar (references V, W) **(Figure 4)**.

Two major clades from Chirundu and one from Magweto, isolated from *Miniopterus, Rhinolophus* and *Hipposideros* spp. of bats also phylogenetically clustered with a strain from Mozambique isolated from *Nycteris thebaica* (reference M). One sequence from Magweto isolated from *Rhinolophus* spp. and two from Mabura cave isolated from *Hipposideros* spp., clustered with a strain of bat astrovirus isolated from *Myotis* spp. in Madagascar (references K, L) **(Figure 4)**. A cluster, comprising two clades from Magweto and one sequence from Chirundu and three sequences from Mabura cave isolated from *Rhinolophus* and *Hipposideros,* were closely related to bat astrovirus strains from *Hipposideros* spp. of Mozambique and Laos (references H, I) **(Figure 4)**. The last small group comprising one clade from Chirundu, four sequences from Magweto, all isolated from *Rhinolophus* spp., and one sequence from Mabura isolated from *Hipposideros* spp. clustered together with bat astrovirus strains from *Rhinolophus* spp. of Laos and Korea (references D, E, F) **(Figure 4)**.

One of the large clusters shows the phylogenetic relationship amongst sequences derived from, *Hipposideros* spp. (Clades HC-5 to –8), these making up the majority of the sequences in this clade and one isolated from *Miniopterus* spp., one from *E. helvum* and one from a *Nycteris* spp.. The bat astroviruses in this clade, all showed a close relation to a bat astrovirus strain isolated from *Hipposideros spp.* of Mozambique (reference C). The largest cluster constituted mainly by Magweto site sequences and some Chirundu sequences, from *Hipposideros* spp. (clades HC-1 to 4) and a few *Rhinolophus* and *Nycteris* spp., clustered with a strain from *Hipposideros* spp. of Mozambique (references A, B) **(Figure 4)**.

## Discussion

The overall AstVs prevalence was respectively 10.0% and 13.7% for Chirundu and Magweto sites, confirming their circulation in bat populations from Zimbabwe. However, compared to studies in other African countries, the observed prevalence was low: Hoarau et *al* and Lebarbenchon et *al* detected AstV in 20-22% of individual bats in Mozambique and Madagascar, respectively (<u>Lebarbenchon et al., 2017</u>; <u>Hoarau et al., 2018</u>). This difference in prevalence of AstV compared to other countries can be attributed to different study designs, longitudinal sampling in our case versus transversal studies in Mozambique and Madagascar (<u>Lebarbenchon et al., 2017</u>; <u>Hoarau et al., 2018</u>). Furthermore, these differences could also result from the studied bat species having varying immunological responses and susceptibilities to different viruses (<u>Hoarau et al., 2023</u>), and the different localities resulting in subtle changes that may impact the infection dynamics.

We also observed difference between the two main sites studied (Chirundu and Magweto). Furthermore, the differences in prevalence between the two main individual populations can be attributed but not limited to differences between habitats, bat population structures and bat species composition of bat communities and also anthropogenic factors. Therefore there is need for a more comprehensive study, covering several years of follow up, to fully establish the factors contributing to the AstV viral dynamics among bat populations at these two sites. At Chirundu site increasing RNA-AstV prevalences were observed from the lactation to the 4-6 months old juvenile seasons with 21.2% of RNA-AstV detection during the latter. Astroviruses have been described to display seasonal variations in prevalence (<u>Lebarbenchon</u> <u>et al., 2017</u>). Drexler et *al* observed different peaks of astrovirus detection associated with different stages of the reproductive season with peaks correlating to maternal aggregations during breeding season and after parturition season due to establishment of a susceptible subpopulation of weaned new-born bats who did not yet have their own adaptive immunity <u>(Lee</u> <u>et al., 2018</u>, <u>Drexler et al., 2011</u>). In our study the prevalence was high during some of the months compared to overall prevalence, which might be related to the effect of reproductive cycle stages on the shedding of viruses, as demonstrated for other viruses. In another study the reproductive season with juveniles and immature individuals showed a very high prevalence of CoV as compared to prevalences in the absence of juveniles and presence of sexually mature bats thus suggesting different stages of the reproduction having effect on viral shedding (<u>Mendenhall et al., 2017</u>; <u>Wacharapluesadee et al., 2018</u>; <u>Cappelle et al., 2021</u>; <u>Chidoti et al.,</u> <u>2022</u>). In this study we observed a similar trend for AstVs for one of the study sites. Furthermore, Mendenhall et *al*, also identified the bat juvenile stage as exhibiting a greater AstVs viral burden than any other stage of the reproductive season (<u>Mendenhall et al., 2017</u>). The range of prevalence observed during the highest peaks coincides with the ≥20% overall AstV detection reported by Hoarau et *al* (<u>Hoarau et al., 2018</u>) during the time of their sampling in February and May. Here we observe a peak during the 4-6 months old juvenile season in Chirundu site which corresponds to a high influx of immunologically immature individuals in the bat population, similar to what was observed in the Chidoti et *al*, for CoV infection (<u>Chidoti</u> <u>et al., 2022</u>).

Juveniles and immature individuals are known to contribute to shedding pulses of viruses as they develop productive infections during the acute phase (<u>Plowright et al., 2016</u>), which coincides with the waning protection of maternal antibodies (<u>Chidoti et al., 2022</u>), thus increasing the susceptibility and rate of infection in the young.

We could not observe a clear trend at Magweto site, which might be related to the absence of samples from December 2020 to March 2021, which correspond to the lactation and weaning periods.

In our study we also investigated co-infection of bats by coronaviruses and astroviruses. We compared the co-infection prevalence with the prevalence of astroviruses and coronaviruses described at each site during the same reproduction cycle stages. The coinfection prevalence showed a similar trend to that observed in the individual viruses, whereby during observed peaks of high AstVs and CoVs detections, the co-infection also peaked at similar phases of the reproduction cycle stages. Thus the co-infection of bats by both CoVs and AstVs is high when there is high infection of bats by either virus.

The overall prevalence of co-infection was 3.5% and 2.6 % in Chirundu and Magweto respectively while the bats from Madagascar and Mozambique respectively showed a AstV-CoV co-infection of 5% (±2.7%) and of 1.7% (±1,1%) (<u>Fischer et al., 2016</u>; <u>Hoarau et al.,</u> <u>2021</u>). Co-infection and recombination of viruses in bats have been reported on several occasions including co-infection of bats by coronaviruses and astroviruses (<u>Fischer et al., 2016</u>; <u>Hoarau et al., 2021, 2023</u>). Bats infected with either CoV or AstV were shown to be more likely co-infected with the respective virus (<u>Seltmann et al., 2017</u>). In our study, no effect of coronavirus infection/shedding on astrovirus infection/shedding was observed.

In the current study we detected bat Astroviruses from *Hipposideridae, Rhinolophidae*, *Pteropodidae, Nycterida*e and *Miniopterida*e families. Astroviruses are known to show no host restriction and are widespread within the Chiroptera order (<u>Chu et al., 2008</u>; <u>Lee et al., 2018</u>). The majority of the astroviruses amplified were from *Hipposideros* and *Rhinolophus* species and these clustered with AstV sequences derived from Mozambique, Madagascar and China of the same families (<u>Lebarbenchon et al., 2017</u>; <u>Hoarau et al., 2018</u>; <u>Lee et al., 2018</u>). The observed trend is due to the two genera being the most dominant species of bats at both sites where sampling was done. Therefore, the probability of higher detection of AstV in these species is expected and evidently observed to be higher than other rare or less dominant species. Active transmission events of astroviruses amongst *Rhinolophidae* and *Hipposideridae* bat species are known to occur (<u>Lee et al., 2018</u>). In this study, a high degree of species-specific tropism, especially in bat astrovirus related clusters was observed with astrovirus strains isolated from *Hipposideros* spp. clustering with each other and similar trends for *Rhinolophus* and *Miniopterus* spp. However, the clusters were not site-specific as sequences from Magweto, Mabura and Chirundu sites formed clusters with each other. We also observed a large phylogenetic cluster which comprised of specific *Hipposideros and Rhinolophus* AstV sub-clades (HC-1 to –8 and Rhi-1 to –4) as well as a few AstV sequences from other bat species (**Figure 4**). Previous studies on astroviruses in bats have also described this family of viruses to be involved in species tropism. However recent studies challenged this assumption and highlighted the fact that AstV cross-species transmissions are more frequent than previously thought (<u>Xiao et al., 2011</u>; <u>Boujon et al., 2017</u>). In our study, AstVs sequences isolated from different bat species of the same site clustered together. This indicates potential cross species transmission of astroviruses within each site, as reported by Xiao et *al* in bats sharing the same habitat (<u>Xiao et al., 2011</u>). We also observed phylogenetic clustering of astroviruses from Chirundu and Magweto sites into same clades showing a great diversity of astroviruses in circulation within the studied bat communities. Novel astroviruses species have been reported in bats hosts (<u>Bergner et al., 2021</u>), and in this study it is evident due to the diverse astroviruses described. The AstV isolated from *E. helvum*, frugivorous bats, clustered together and showed close phylogenetic relation to *BAstV* isolated from *Rousettus* spp. bats in China and *Taphozous* spp. bats in Laos. One strain isolated from *E. helvum* clustered with *Hipposideros* bat astroviruses sequences isolated in Chirundu site. This could mean that frugivorous bats and insectivorous bats can be infected by the same type of AstVs strain thus indicative of the multi-host spectrum for the Astroviridae family. Detection of AstVs in the frugivorous bats was low compared to insectivorous bats, and this can be attributed to the insectivorous bats ability to adapt and harbour more astroviruses than frugivorous bats (<u>Xiao et al., 2011</u>).

Two strains, Mag-292 *H. Caffer*, Mag-573 *Rhinolophus* spp. clustered within the *HuAstv* type 1 to 6, that are known to cause infection and diarrhoea in children and infants (<u>Bosch et al., 2014</u>). Six of our strains isolated from *E. helvum* and one from *Rhinolophus* spp. formed sister clades to *HuAstV* HMO, associated with severe extra-intestinal illnesses in humans (<u>Wohlgemuth et al., 2019</u>) and VA-1-4 strains. Another AstV strain detected in *Hipposideros* bat species clustered with *HuAstv* MLB-1, –2 and –3 which are all known to cause gastro-intestinal and neurological diseases with mild-severe symptoms in humans (<u>Kapoor et</u> <u>al., 2009</u>). This phylogenetic clustering could suggest that *HuAstV* and *BAstV* may have a shared common ancestor. However, with limited sequence data the evolutionary history is merely a speculation that needs to be further investigated through full genome analysis. Another hypothesis for the observed relation is that, since bats are known to be important reservoir hosts of astroviruses with great diversity, they could be ancestors of the *HuAstV* due to the high prevalence of AstV detected in bats. *HuAstV* transmission occurs via the faecal-oral route and contaminated food or water (<u>Bosch et al., 2014</u>), therefore it is unlikely that anthroponosis may have occurred. However, this observed relation could be due to a rare event or mutation, or a new strain of bat astrovirus that has not been described yet. Further full genomic data as well as epidemiological data on these bat colonies are needed to reach a full conclusion.As novel strains and host species continue to be discovered, understanding AstV transmission and their zoonotic potential is essential. Dissemination of *HuAstV* and AstV from zoonotic reservoir hosts remain a significant threat to public health (<u>Roach & Langlois, 2021</u>).

## Conclusion

This study offered important contributions to the understanding of virus diversity in an under-sampled region of the world, and the sequences contribute to a better picture of co-evolution and transmission among bat species. Multiple sampling points as in the study further give a more comprehensive picture of how viruses vary over space and time and host species. Astroviruses have a high genetic diversity, multiple mechanism of generating additional diversity, and infect a wide range of host species therefore understanding their prevalence and infection dynamics in their wildlife hosts can help to predict or prevent the emergence of novel astrovirus strains into domestic animals and human populations.

## Acknowledgments

We thank Billy Butete for his field assistance. The authors gratefully acknowledge and thank the International Atomic Energy Agency for making available the sequencing services. We thank the Research Council of Zimbabwe for approving this study (research registration certificate N° 03006) and the Hurungwe Rural District council for their assistance and facilitation. We thank the Animal Research Ethics Committee of Zimbabwe for their approval (ref number 002/2017). This work was conducted within the framework of the Research Platform “Production and Conservation in Partnership” (RP–PCP).

The Preprint version 5 of this article has been peer-reviewed and recommended by Peer Community in PCI infection: DOI: <u>10.24072/pci.infections.100085</u> (James T., 2023).

## Author’s contribution: Conceptualization

VC, MA, HdN, MB, LG and FL.; **Methodology**: VC, MA, LG, HdN, DP, EG, GM, EM and FL.; **Software:** VC, MA, HdN and FL; **Validation**: VC, MA HdN, JC, and FL; **Formal Analysis**: VC, HdN, JC, and FL.; **Investigation**: VC, VP, NC, GM, and FL.; **Resources**: DP, GM, MP, MB, HdN, LG, and FL.; **Data Curation**: VC, HdN, and FL.; **Writing – Original Draft Preparation**: VC and FL; **Writing – Review & Editing**: HdN, MA, JC, MB, LG, VP, EG, DM, DP, EM, GM and FL; **Supervision**: HdN, DP, GM and FL.; **Project Administration**: LG, MB, HdN and FL; **Funding Acquisition**: MB, HdN, LG and FL.

**Data, scripts, code, and supplementary information availability**

***Sampling Data, Astrovirus sequences, Graphs and Figures***

https://doi.org/10.5281/zenodo.8399450

https://doi.org/10.5281/zenodo.7849008

## GenBank Accession Numbers

The AstVs sequences have been deposited to the GenBank under the following numbers: OQ271049 – OQ271203

The Cytochrome B sequences have been deposited to the GenBank under the following numbers: OM487705-OM488020

## Statistical analysis

The script used for the GLMM analysis is available at: https://doi.org/10.5281/zenodo.7847934

***Supplementary data***: https://doi.org/10.5281/zenodo.8399361

## Reference**s

1. Amman BR, Nyakarahuka L, McElroy AK, Dodd KA, Sealy TK, Schuh AJ, Shoemaker TR, Balinandi S, Atimnedi P, Kaboyo W, Nichol ST, Towner JS (2014) Marburgvirus resurgence in Kitaka mine bat population after extermination attempts, Uganda. Emerging Infectious Diseases, 20, 1761–1764. 10.3201/eid2010.140696

2. Baker KS, Todd S, Marsh GA, Crameri G, Barr J, Kamins AO, Peel AJ, Yu M, Hayman DTS, Nadjm B, Mtove G, Amos B, Reyburn H, Nyarko E, Suu-Ire R, Murcia PR, Cunningham AA, Wood JLN, Wang L-FL-F (2013) Novel, potentially zoonotic paramyxoviruses from the African straw-colored fruit bat Eidolon helvum. Journal of virology, 87, 1348–58. 10.1128/JVI.01202-12

3. Bergner LM, Mollentze N, Orton RJ, Tello C, Broos A, Biek R, Streicker DG (2021) Characterizing and Evaluating the Zoonotic Potential of Novel Viruses Discovered in Vampire Bats. Viruses, 13, 252. 10.3390/v13020252

4. Bosch A, Pintó RM, Guix S (2014) Human astroviruses. Clinical Microbiology Reviews, 27, 1048–1074. 10.1128/CMR.00013-14

5. Boujon CL, Koch MC, Seuberlich T (2017) The Expanding Field of Mammalian Astroviruses: Opportunities and Challenges in Clinical Virology. Advances in Virus Research, 99, 109–137. 10.1016/bs.aivir.2017.07.002

6. Bourgarel M, Pfukenyi DM, Boué V, Talignani L, Chiweshe N, Diop F, Caron A, Matope G, Missé D, Liégeois F (2018) Circulation of Alphacoronavirus, Betacoronavirus and Paramyxovirus in Hipposideros bat species in Zimbabwe. Infection, Genetics and Evolution, 58, 253–257. 10.1016/j.meegid.2018.01.007

7. Cappelle J, Furey N, Hoem T, Ou TP, Lim T, Hul V, Heng O, Chevalier V, Dussart P, Duong V (2021) Longitudinal monitoring in Cambodia suggests higher circulation of alpha and betacoronaviruses in juvenile and immature bats of three species. Scientific Reports, 11, 1–11. 10.1038/s41598-021-03169-zz

8. Chidoti V, De Nys H, Pinarello V, Mashura G, Misse M, Guerrini L, Pfukenyi D, Cappelle J, Chiweshe N, Ayouba A, Matope G, Peeters M, Gori E, Bourgarel M, Liegeois F (2022) Longitudinal survey of Coronavirus circulation and diversity in insectivorous bat colonies in Zimbabwe. Viruses, 14, 781. 10.3390/v14040781

9. Chu DKW, Poon LLM, Guan Y, Peiris JSM (2008) Novel Astroviruses in Insectivorous Bats. Journal of Virology, 82, 9107–9114. 10.1128/JVI.00857-08

10. Cortez V, Meliopoulos VA, Karlsson EA, Hargest V, Johnson C, Schultz-Cherry S (2017) Astrovirus Biology and Pathogenesis. Annual Review of Virology, 4, 327–348. 10.1146/annurev-virology-101416-041742

11. Donato C, Vijaykrishna D (2017) The broad host range and genetic diversity of mammalian and avian astroviruses. Viruses, 9, 1–18. 10.3390/v9050102

12. Drexler JF, Corman VM, Wegner T, Tateno AF, Zerbinati RM, Gloza-Rausch F, Seebens A, Müller MA, Drosten C (2011) Amplification of emerging viruses in a bat colony. Emerging Infectious Diseases, 17, 449–456. 10.3201/eid1703.100526

13. Dufkova L, Straková P, Širmarová J, Salát J, Moutelíková R, Chrudimský T, Bartonička T, Nowotny N, Růžek D (2015) Detection of Diverse Novel Bat Astrovirus Sequences in the Czech Republic. Vector-Borne and Zoonotic Diseases, 15, 518–521. 10.1089/vbz.2015.1813

14. Fischer K, dos Reis VP, Balkema-Buschmann A (2017) Bat astroviruses: Towards understanding the transmission dynamics of a neglected virus family. Viruses, 9, 8–16. 10.3390/v9020034

15. Fischer K, Zeus V, Kwasnitschka L, Kerth G, Haase M, Groschup MH, Balkema-Buschmann A (2016) Insectivorous bats carry host specific astroviruses and coronaviruses across different regions in Germany. Infection, Genetics and Evolution, 37, 108–116. 10.1016/j.meegid.2015.11.010

16. Frémond M-L, Pérot P, Muth E, Cros G, Dumarest M, Mahlaoui N, Seilhean D, Desguerre I, Hébert C, Corre-Catelin N, Neven B, Lecuit M, Blanche S, Picard C, Eloit M (2015) Next-Generation Sequencing for Diagnosis and Tailored Therapy: A Case Report of Astrovirus-Associated Progressive Encephalitis. Journal of the Pediatric Infectious Diseases Society, 4, e53–e57. 10.1093/jpids/piv040

17. Guindon S, Dufayard JF, Lefort V, Anisimova M, Hordijk W, Gascuel O (2010) New algorithms and methods to estimate maximum-likelihood phylogenies: Assessing the performance of PhyML 3.0. Systematic Biology, 59, 307–321. 10.1093/sysbio/syq010

18. Hoarau AOG, Goodman SM, Halabi D Al, Ramasindrazana B, Lagadec E, Minter G Le, Köster M, Santos A Dos, Schoeman MC, Gudo ES, Mavingui P, Lebarbenchon C (2021) Investigation of astrovirus, coronavirus and paramyxovirus co – infections in bats in the western Indian Ocean. Virology Journal, 1–8. 10.1186/s12985-021-01673-2

19. Hoarau AOG, Köster M, Dietrich M, Le Minter G, Joffrin L, Ramanantsalama R V., Mavingui P, Lebarbenchon C (2023) Synchronicity of viral shedding in molossid bat maternity colonies. Epidemiology and Infection, 151. 10.1017/S0950268823000171

20. Hoarau F, Minter G Le, Joffrin L, Schoeman MC, Lagadec E, Ramasindrazana B, Santos A Dos, Goodman SM, Gudo ES, Mavingui P, Lebarbenchon C (2018) Bat Astrovirus in Mozambique. Virology Journal (2018), **6**, 1–5. 10.1186/s12985-018-1011-x

21. ICTV (2020) ICTV Positive Sense RNA Viruses: Astroviridae. VIRUS Taxon.

22. Janowski AB, Klein RS, Wang D (2019) Differential In Vitro Infection of Neural Cells by Astroviruses (DE Griffin, Ed,). mBio, 10. 10.1128/mBio.01455-19

23. Johnson CK, Hitchens PL, Pandit PS, Rushmore J, Evans TS, Young CCW, Doyle MM, Johnson CK (2020) Global shifts in mammalian population trends reveal key predictors of virus spillover risk. Proc. R. Soc. B, 287. 10.1098/rspb.2019.2736

24. Kapoor A, Li L, Victoria J, Oderinde B, Mason C, Pandey P, Zaidi SZ, Delwart E (2009) Multiple novel astrovirus species in human stool. Journal of General Virology, 90, 2965– 2972. 10.1099/vir.0.014449-0

25. Kocher TD, Thomas WK, Meyer A, Edwards S V., Paabo S, Villablanca FX, Wilson AC (1989) Dynamics of mitochondrial DNA evolution in animals: Amplification and sequencing with conserved primers. Proceedings of the National Academy of Sciences of the United States of America, 86, 6196–6200. 10.1073/pnas.86.16.6196

26. Kumar S, Stecher G, Tamura K (2016) MEGA7: Molecular Evolutionary Genetics Analysis Version 7.0 for Bigger Datasets. Molecular Biology and Evolution, 33, 1870–1874. 10.1093/molbev/msw054

27. Lacroix A, Duong V, Hul V, San S, Davun H, Omaliss K, Chea S, Hassanin A, Theppangna W, Silithammavong S, Khammavong K, Singhalath S, Afelt A, Greatorex Z, Fine AE, Goldstein T, Olson S, Joly DO, Keatts L, Dussart P, Frutos R, Buchy P (2017) Diversity of bat astroviruses in Lao PDR and Cambodia. Infection, Genetics and Evolution, 47, 41–50. 10.1016/j.meegid.2016.11.013

28. Larkin MA, Blackshields G, Brown NP, Chenna R, Mcgettigan PA, McWilliam H, Valentin F, Wallace IM, Wilm A, Lopez R, Thompson JD, Gibson TJ, Higgins DG (2007) Clustal W and Clustal X version 2.0. Bioinformatics, 23, 2947–2948. 10.1093/bioinformatics/btm404

29. Lebarbenchon C, Ramasindrazana B, Joffrin L, Bos S, Lagadec E, Le Minter G, Gomard Y, Tortosa P, Wilkinson DA, Goodman SM, Mavingui P (2017) Astroviruses in bats, Madagascar. Emerging Microbes & Infections, 6, 1–3. 10.1038/emi.2017.47

30. Lee S-Y, Son K-D, Yong-Sik K, Wang S-J, Kim Y-K, Jheong W-H, Oem J-K (2018) Genetic diversity and phylogenetic analysis of newly discovered bat astroviruses in Korea. Archives of Virology, 163, 3065–3072. 10.1007/s00705-018-3992-6

31. Lemoine F, Wilkinson E, Correia D, Oliveira D, Gascuel O (2018) Renewing Felsenstein’s Phylogenetic Bootstrap in the Era of Big Data. 10.1038/s41586-018-0043-0.

32. Mendenhall IH, Skiles MM, Neves ES, Borthwick SA, Low DHW, Liang B, Lee BPYH, Su YCF, Smith GJD (2017) Influence of age and body condition on astrovirus infection of bats in Singapore: An evolutionary and epidemiological analysis. One Health, 4, 27–33. 10.1016/j.onehlt.2017.10.001

33. Milne I, Lindner D, Bayer M, Husmeier D, Mcguire G, Marshall DF, Wright F (2009) TOPALi v2: A rich graphical interface for evolutionary analyses of multiple alignments on HPC clusters and multi-core desktops. Bioinformatics, 25, 126–127. 10.1093/bioinformatics/btn575

34. Monadjem A, Taylor PJ, Cotterill FPD, Schoeman MC (2010) Bats of southern and central Africa: A biogeographic and taxonomic synthesis. Wits University Press, Johannesburg. 10.1644/12-MAMM-R-184.1

35. Plowright RK, Peel AJ, Streicker DG, Gilbert AT, McCallum H, Wood J, Baker ML, Restif O (2016) Transmission or Within-Host Dynamics Driving Pulses of Zoonotic Viruses in Reservoir–Host Populations (J V. Remais, Ed,). PLOS Neglected Tropical Diseases, 10, e0004796. 10.1371/journal.pntd.0004796

36. Roach SN, Langlois RA (2021) Intra and Cross-Species Transmission of Astroviruses. Viruses, 13, 1127. 10.3390/v13061127

37. Rougeron V, Suquet E, Maganga GD, Jiolle D, Mombo IM, Bourgarel M, Motsch P, Arnathau C, Durand P, Drexler F, Drosten C, Renaud F, Prugnolle F, Leroy EM (2016) Characterization and phylogenetic analysis of new bat astroviruses detected in Gabon, Central Africa. Acta virologica, 60, 386–392. 10.4149/av_2016_04_386

38. Seltmann A, Corman VM, Rasche A, Drosten C, Czirják GÁ, Bernard H, Struebig MJ, Voigt CC (2017) Seasonal Fluctuations of Astrovirus, But Not Coronavirus Shedding in Bats Inhabiting Human-Modified Tropical Forests. EcoHealth, 14, 272–284. 10.1007/s10393-017-1245-x

39. El Taweel A, Kandeil A, Barakat A, Rabiee OA, Kayali G, Ali MA (2020) Diversity of astroviruses circulating in humans, bats, and wild birds in Egypt. Viruses, 12, 1–12. 10.3390/v12050485

40. Vu DL, Cordey S, Brito F, Kaiser L (2016) Novel human astroviruses: Novel human diseases? Journal of Clinical Virology, 82, 56–63. 10.1016/j.jcv.2016.07.004

41. Wacharapluesadee S, Duengkae P, Chaiyes A, Kaewpom T, Rodpan A, Yingsakmongkon S, Petcharat S, Phengsakul P, Maneeorn P, Hemachudha T (2018) Longitudinal study of age-specific pattern of coronavirus infection in Lyle’s flying fox (Pteropus lylei) in Thailand. Virology Journal, 15, 1–10. 10.1186/s12985-018-0950-6

42. Waruhiu C, Ommeh S, Obanda V, Agwanda B, Gakuya F, Ge XY, Yang X Lou, Wu LJ, Zohaib A, Hu B, Shi ZL (2017) Molecular detection of viruses in Kenyan bats and discovery of novel astroviruses, caliciviruses and rotaviruses. Virologica Sinica, 32, 101–114. 10.1007/s12250-016-3930-2

43. Wilson E. (1927) Probable Inference, the Law of Succession, and Statistical Inference. Journal of the American Statistical Association, 22, 209–212. 10.2307/2276774

44. Wohlgemuth N, Honce R, Schultz-Cherry S (2019) Astrovirus evolution and emergence. Infection, Genetics and Evolution, 69, 30–37. 10.1016/j.meegid.2019.01.009

45. Wu H, Pang R, Cheng T, Xue L, Zeng H, Lei T, Chen M, Wu S, Ding Y, Zhang J, Shi M, Wu Q (2020) Abundant and Diverse RNA Viruses in Insects Revealed by RNA-Seq Analysis: Ecological and Evolutionary Implications (IM Cristea, Ed,). mSystems, 5. 10.1128/mSystems.00039-20

46. Xiao J, Li J, Hu G, Chen Z, Wu Y, Chen Y, Chen Z, Liao Y, Zhou J, Ke X, Ma L, Liu S, Zhou J, Dai Y, Chen H, Yu S, Chen Q (2011) Isolation and phylogenetic characterization of bat astroviruses in southern China. Archives of Virology, 156, 1415–1423. 10.1007/s00705-011-1011-2

